# The non-kinase function of CDK6 is a key driver of acquired resistance to CDK4/6 inhibitors in Estrogen Receptor-positive breast cancer

**DOI:** 10.1101/2025.11.11.687883

**Authors:** S M Nashir Udden, Baishan Jiang, Kamal Pandey, Cheryl M. Lewis, Sunati Sahoo, Noelle S. Williams, Prasanna G. Alluri

## Abstract

The therapeutic efficacy of CDK4/6 inhibitors in ER+ breast cancer (BC) patients is presumed to arise from inhibition of the kinase function of CDK4 and 6 proteins. Despite their initial efficacy, development of acquired resistance to CDK4/6 inhibitors in metastatic ER+ BC patients is nearly universal, commonly through compensatory overexpression of CDK6. Here, we show that primary ER+ tumors show high CDK4 but very low to undetectable expression of CDK6, which suggests that the therapeutic efficacy of CDK4/6 inhibitors in ER+ BC patients is largely driven by inhibition of CDK4, rather than CDK6. Furthermore, we show that overexpression of CDK6 confers acquired resistance to CDK4/6 inhibitors, largely, through a kinase-independent function. Our findings challenge the dogma that CDK4/6-mediated phosphorylation of retinoblastoma protein (pRB) is necessary to ER+ BC cells to progress from G1 to S phase of cell cycle. Consequently, by taking advantage of the non-kinase function of CDK6, ER+ BC cells acquire independence from the kinase function of CDK4 and 6 for cell cycle progression. These findings highlight the limitations of the current standard-of-care treatments, which focus on merely inhibiting the kinase function of CDK4 and 6, and uncover the non-kinase function of CDK6 as a new targetable vulnerability to overcome acquired resistance to CDK4/6 inhibitors in ER+ BC.

---

CDK4/6 inhibitors such as palbociclib, when combined with an endocrine therapy, substantially improve progression-free survival in patients with metastatic, ER+ BC (*1*). However, development of acquired resistance to CDK4/6 inhibitors in this patient population is nearly universal (*1, 2*). Overexpression of CDK6 is a common acquired mechanism of resistance to CDK4/6 inhibitors in metastatic ER+ BC patients (*3-5*). Diverse genetic mechanisms such as loss of *FAT1, PTEN and ARIAD1* as well as genomic amplification of CDK6 drive overexpression of CDK6 in metastatic ER+ breast tumors following treatment with CDK4/6 inhibitors (*5*).

To probe the dependence of treatment naïve, ER+ BC cells on CDK4 and CDK6 for growth, we measured the expression levels of CDK4 and CDK6 in a panel of hormone receptor-positive (HR+), Human Epidermal Growth Factor Receptor 2-negative (HER2-) BC cell lines by Western blot analysis. Surprisingly, none of the cells tested showed detectable expression of CDK6, although they showed high expression of CDK4 **(Fig. 1A)**. In contrast, both CDK4 and CDK6 protein levels were high in a panel of Triple Negative Breast Cancer (TNBC) cell lines **(Fig. 1A)**. These findings were corroborated in an institutional cohort of primary tumors from 33 HR+, HER2-BC patients in which 28/33 patients (85%) showed detectable CDK4 expression and 0/33 patients showed detectable CDK6 expression **(Fig. B)**. To further validate these findings, we probed RNA-seq data from a panel of HR+, HER2- and TNBC patient-derived xenografts (PDXs) **(Table ST1)** (*6*). Expression of CDK4 did not vary significantly between these PDXs **(Fig. 1C)**. However, in nearly all cases, TNBC PDXs exhibited high expression of CDK6 while HR+, HER2-PDXs universally showed very low mRNA expression **(Fig. 1C)**. Thus, several independent lines of evidence show that HR+, HER2-BC cells and tumors express low to undetectable levels of CDK6 mRNA and protein. These findings suggest that the activity of CDK4/6 inhibitors in ER+ breast tumors is primarily driven by the inhibition of CDK4, rather than CDK6, since the expression levels of CDK6 in these tumors are very low or undetectable. Consequently, CDK6 inhibition in ER+ BC patients (who have not been previously treated with a CDK4/6 inhibitor) may contribute to toxicity without necessarily affording a therapeutic benefit since these tumors exhibit very low or undetectable CDK6. In fact, many hematological adverse effects such as altered erythroid differentiation and neutropenia are attributable to inhibition of CDK6, but not CDK4 function (*7*).

**Figure 1.**
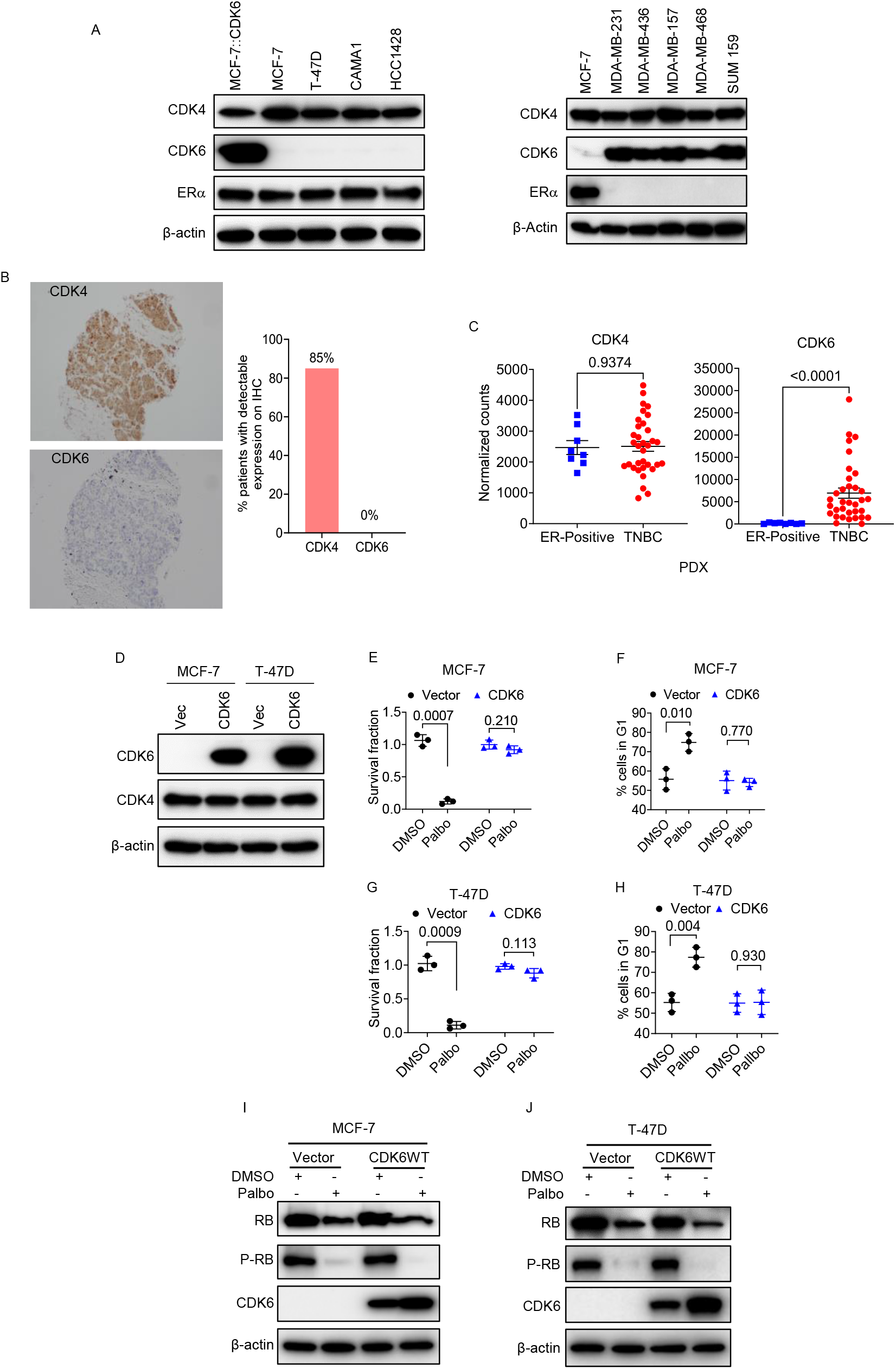
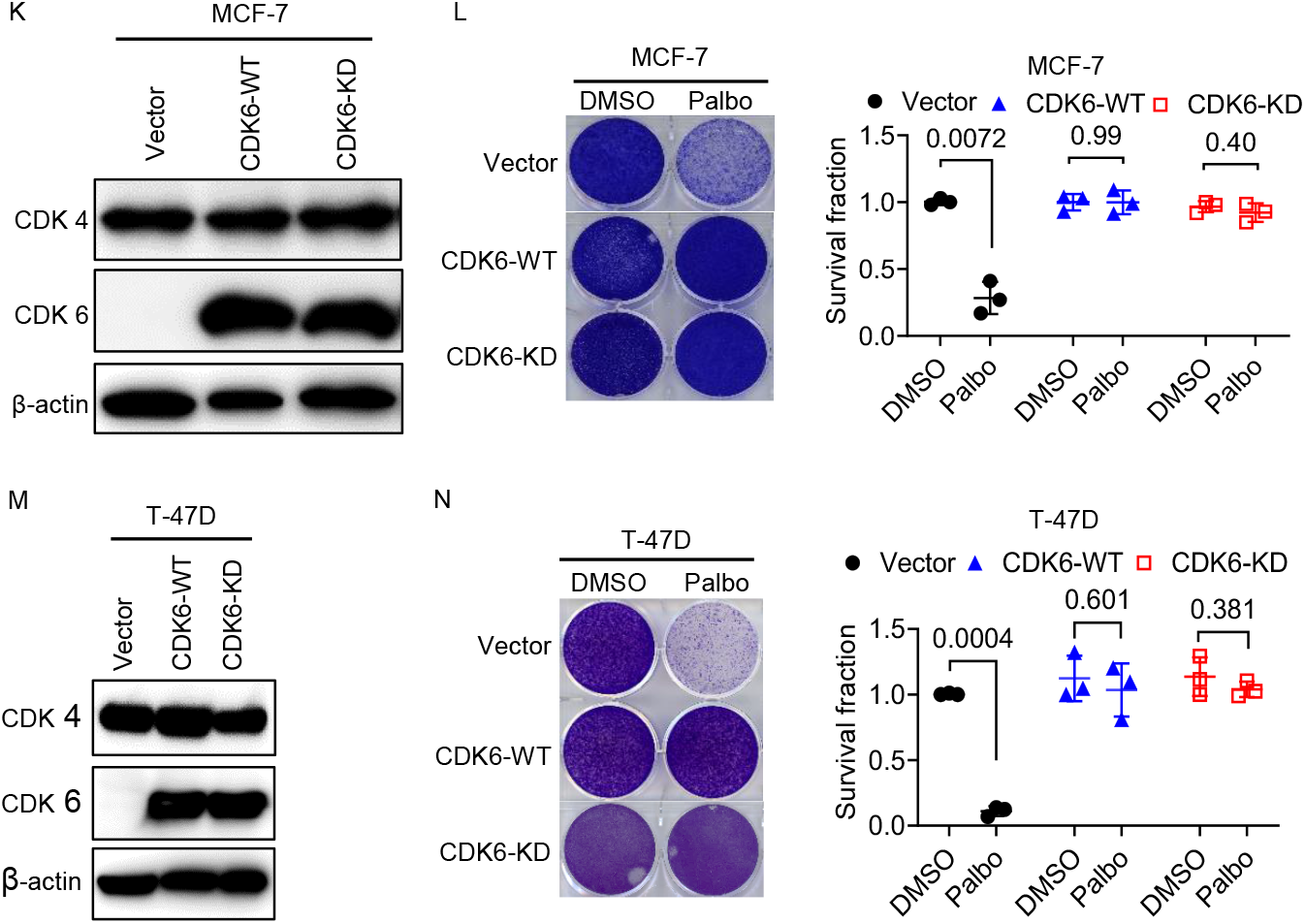
The non-kinase function of CDK6 is a key driver of resistance to CDK4/6 inhibitors in ER+ breast cancer cells. A: CDK4 and 6 expression by Western blot analysis in a panel of ER+ breast cancer cell lines (left) and Triple Negative Breast Cancer (TNBC) cell lines (right). MCF-7 cells ectopically overexpressing CDK6 were used as a positive control. B: Representative immunohistochemical staining showing high CDK4 and no detectable CDK6 in an institutional cohort of 33 primary ER+ breast tumors. 28/33 tumors in this cohort had detectable CDK4 expression and 0/33 had detectable CDK6 expression. C: RNA-seq data from a panel of hormone receptor-positive (HR+), Human Epidermal Growth Factor Receptor 2 (HER-2)-negative (n=8) or TNBC patient-derived xenografts (PDXs) (n=34) showing similar expression of CDK4 in both sub-types but high expression of CDK6 in most TNBC PDXs, and very low expression of CDK6 in HR+, HER-2-negative PDXs. D: Expression of CDK4 and CDK6 by Western blot analysis in MCF-7 and T-47D cells stably transfected with an empty vector or a CDK6 expression vector. E&G: MCF-7 and T-47D cells stably transfected with an empty vector or CDK6 expression vector were treated with vehicle or palbociclib (1 μM) in biological triplicates and cell survival was quantified 10 days later by crystal violet staining. F&H: MCF-7 and T-47D cells stably transfected with an empty vector or CDK6 expression vector were treated with vehicle or palbociclib (1 μM) for 24 hours in biological triplicates and proportion of cells in G1 were quantified by flow cytometry. I&J: MCF-7 or T-47 cells stable transfected with an empty vector or CDK6 expression vector were treated with vehicle or palbociclib (1 μM) for 24 hours and the expression levels of total retinoblastoma protein and phospho-retinoblastoma proteins were analyzed by Western blotting. K-N: MCF-7 or T-47D cells stable transfected with an empty, CDK6-WT or CDK6-KD expression vectors were treated with vehicle or palbociclib (1 μM) in biological triplicates and cell survival was quantified 10 days later by crystal violet staining. WT= wild type; KD= kinase dead. Paired comparisons between groups were made using a two-tailed Student’s t-test. A p value of less than 0.05 was considered statistically significant.

Acquired resistance to CDK4/6 inhibitors in ER+ BC patients is commonly driven by compensatory overexpression of CDK6 (rather than CDK4) (*3-5*), although ER+ tumors naïve to CDK4/6 inhibitor treatment show undetectable expression of CDK6. These findings suggest that CDK6 overexpression may confer a survival advantage (over CDK4 overexpression) to ER+ BC cells in the face of CDK4/6 inhibitor treatment. To recapitulate these findings in a preclinical model, we generated ER+ BC cell lines with stable ectopic overexpression of CDK6 (**Fig. 1D)**. These cells exhibited remarkable resistance to palbociclib while the corresponding parental cells transfected with empty vector were highly sensitive to the drug **(Fig. 1E&1G)**. While palbociclib treatment (1 μM) for 24 hours caused significant enrichment of cells in G1 in parental MCF-7 and T-47D cells transfected with an empty vector (due to drug-induced G1 arrest), overexpression of CDK6 blunted this effect and cells evaded palbociclib-induced G1 arrest **(Fig. 1F&1H)**.

Next, we asked if palbociclib resistance in CDK6-overexpressing ER+ BC cells is driven by incomplete inhibition of pRB phosphorylation due to high expression levels of CDK6. Treatment of MCF-7 and T-47D cells overexpressing CDK6 with palbociclib (1 μM) for 24 hours completely blocked pRB phosphorylation in both the cell lines **(Fig. 1I&1J)**, although these cells were completely resistant to treatment with same concentration of the drug **(Fig. 1E&1G)**. These findings strongly suggest that CDK6-driven resistance to palbociclib is not due to incomplete inhibition of pRB phosphorylation but likely driven by a non-canonical function of CDK6.

To determine whether the non-kinase function of CDK6 is a driver of palbociclib resistance, we expressed a previously characterized kinase-dead (KD) mutant (K43M) of CDK6 (*8*) (referred henceforth as CDK6-KD) in MCF-7 and T-47D cells (**Fig. 1K & Fig. S1A)**. The expression levels of CDK6-WT and CDK6-KD were comparable in these cells. An *in vitro* kinase assay demonstrated that the CDK6 protein immunoprecipitated from MCF-7 cells stably expressing the wild type protein phosphorylated recombinant retinoblastoma protein while immunoprecipitation of CDK6 from MCF-7 cells stably expressing CDK6-KD did not **(Fig. S1)**. Despite lack of demonstrable kinase activity, expression of CDK6-KD in both MCF-7 and T-47D cells conferred resistance to palbociclib (1μM) *in vitro* **(Fig. 1L&N)**. Thus, expression of CDK6-KD was sufficient to confer palbociclib resistance in both MCF-7 and T-47D cells, thereby definitively establishing that the non-kinase function of CDK6 is the primary driver of palbociclib resistance.

Supported by our findings, which show that expression of the kinase-dead mutant of CDK6 was sufficient to confer palbociclib resistance, we hypothesized that eliminating both the kinase and non-kinase functions of CDK6 (rather than mere inhibition of CDK6 kinase function achieved with existing CDK4/6 inhibitors) is necessary to inhibit the growth of BC cells that have developed palbociclib resistance due to CDK6 overexpression **(Fig. 2A)**. To test this hypothesis, we employed a CDK4/6 selective degrader, BSJ-1-184 (referred to as CDK4/6-D from here forth), a PROteolysis TArgeting Chimera (PROTAC) that causes selective degradation of CDK6 and 4 **(Fig. 2A)** (*9*). To determine the specificity of target degradation of CDK4/6-D, we treated Jurkat cells, which express high endogenous levels of CDK4 and 6, with 250 nM CDK4/6D for 6 hours and assessed proteome-wide selectivity of CDK4/6-D in an unbiased fashion. Remarkably, CDK6 was the most significantly depleted protein and CDK4 was among the top 10 depleted proteins across the whole proteome **(Fig, S3)**. Consistent with these results, treatment of two independent ER+ cell lines (MCF-7 and T-47D cells) overexpressing CDK6 with CDK4/6-D (1μM) for 24 hours resulted in significant depletion of both CDK6 and CDK4 proteins **(Fig. 2B&C)**. Furthermore, CDK4/6-D (1μM) inhibited the growth of both MCF-7 and T-47D cells expressing CDK6-WT and CDK6-KD, although these cells were completely resistant to palbociclib (1μM) **(Fig. 2D&E)**.

**Figure 2.**
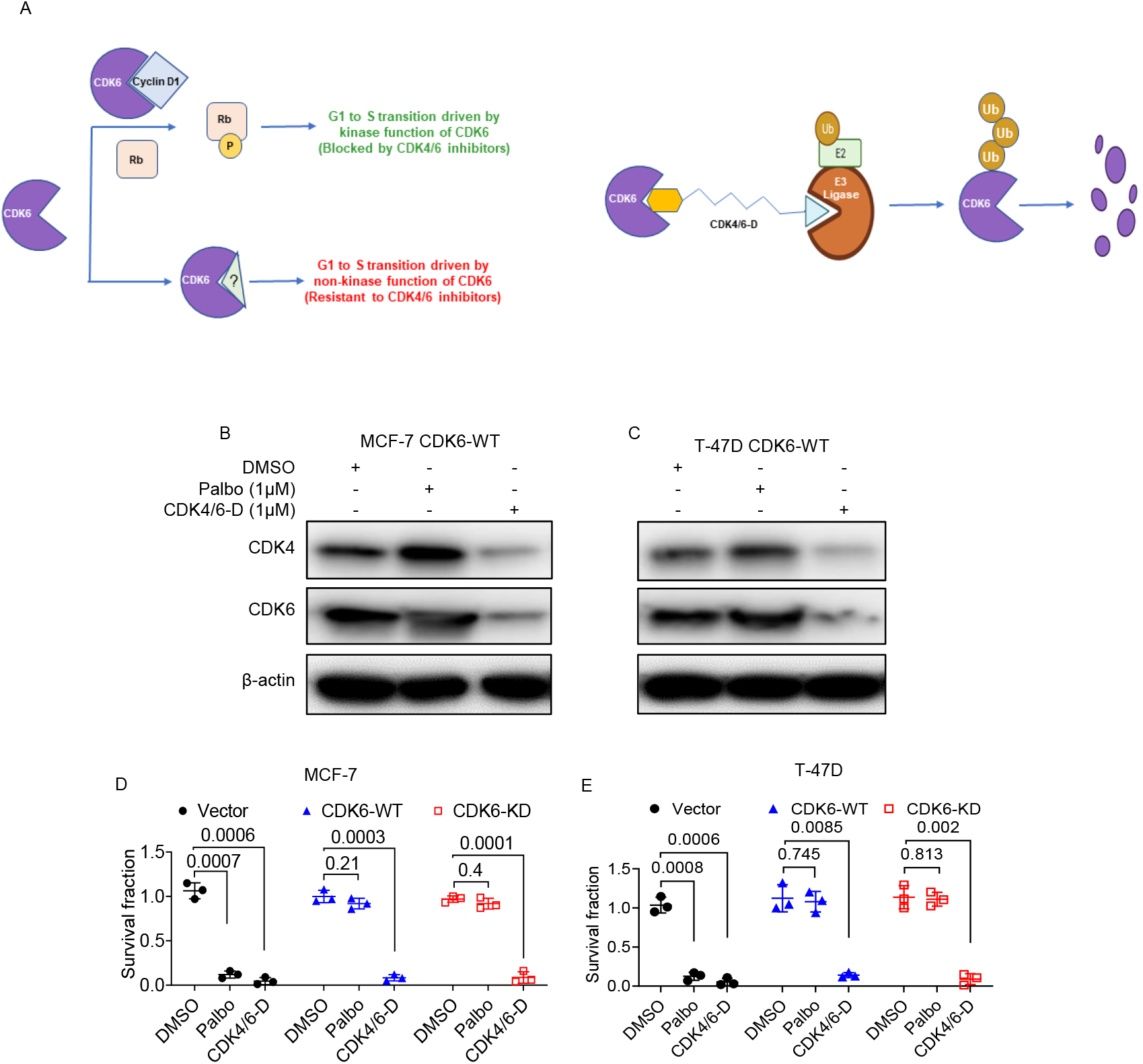
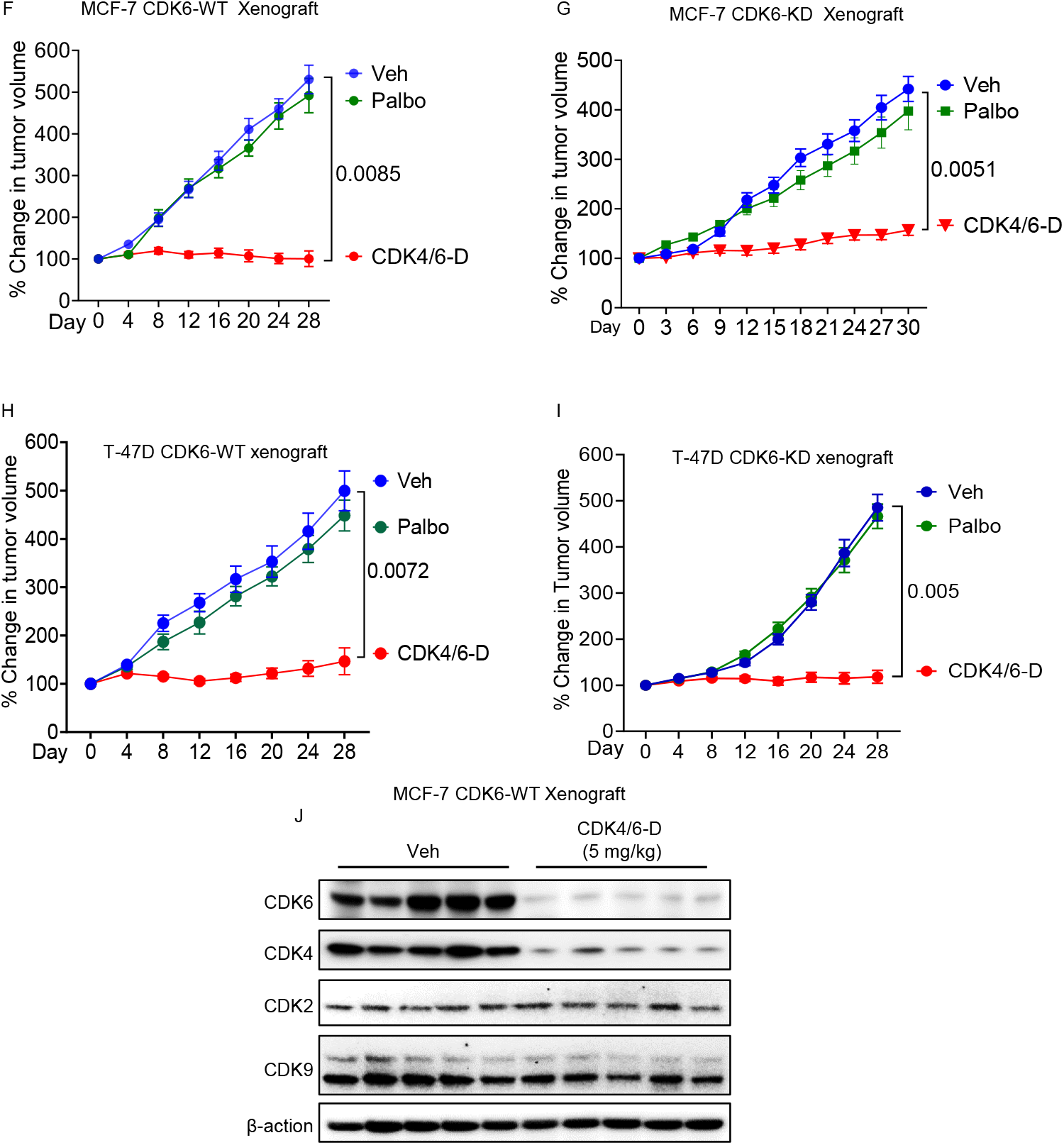
Targeted degradation of CDK6 (and 4) inhibits the growth CDK4/6 inhibitor-resistant xenografts. A: Schematic depiction of dual roles of CDK6 in promoting tumor growth through both kinase-dependent and independent functions (Left). Schematic depiction of CDK4/6-D, a proteolysis targeting chimera (PROTAC), which induces targeted degradation of CDK6 (and 4). B-C: MCF-7 and T-47D cells stably transfected with a CDK6-WT expression vector were treated with vehicle, palbociclib (1 μM) or CDK4/6-D (1 μM) for 24 hours and expression levels of CDK4 and 6 measured by Western blotting. D-E: MCF-7 and T-47D cells stably transfected with an empty, CDK6-WT or CDK6-KD expression vectors were treated with vehicle, palbociclib (1 μM) or CDK4/6-D (1 μM) in biological triplicates and cell survival was quantified 10 days later by crystal violet staining. F-I: MCF-7 CDK6-WT (F) and MCF-7 CDK6-KD (G), T-47D CDK6-WT (H) and T-47D CDK6-KD (I) xenograft-bearing mice (n=10 tumors per treatment group) were treated with vehicle, palbociclib (100 mg/kg daily via oral gavage), or CDK4/6-D (5 mg/kg daily via intraperitoneal injection) for 4 weeks and tumor volumes were measured every 3-4 days. For *in vitro* studies, statistical significance was evaluated using an unpaired, two-tailed t-test. For *in vivo* tumor studies, ANOVA followed by Dunnett’s test was used to adjust for multiple comparisons between the experimental arms and the control arm. J: Representative tumors (n=5) collected from F were subjected to Western blotting and expression levels of CDK2, CDK4, CDK6 and CDK9 were visualized. WT= wild type; KD= kinase dead.

To facilitate *in vivo* evaluation of CDK4/6-D, we carried out pharmacokinetic (PK) studies after administering a single dose of CDK4/6-D (50 mg/kg) by intraperitoneal (i.p.) injection and achieved excellent plasma exposure **(Fig. S3)**. To evaluate *in vivo* efficacy, we carried out therapeutic studies at a drug dose of 5 mg/kg (daily i.p. injection). Mice did not show evidence of toxicity at this dose as assessed by body weight loss or through biochemical tests to assess organ function **(Fig. S4)**. For xenograft studies, we injected MCF-7 cells engineered to express CDK6-WT into 6–8-week-old, ovariectomized female athymic, nude mice following subcutaneous implantation of 0.17 mg β-estradiol pellet (2-week release). When tumors reached ∼100-150 mm^3^, mice were treated with vehicle, palbociclib (100 mg/kg daily by oral gavage) or CDK4/6-D (5 mg/kg daily by i.p. injection) for 4 weeks. As expected, xenografts overexpressing CDK6-WT were completely resistant to palbociclib **(Fig. 2F)**. Remarkably, these xenografts achieved complete growth inhibition when treated with CDK4/6-D **(Fig. 2F)**.

To definitively establish the role of CDK6-KD in conferring palbociclib resistance *in vivo*, we generated xenografts from MCF-7 CDK6-KD cells and treated them with vehicle, palbociclib or CDK4/6-D. CDK6-KD xenografts exhibited remarkable resistance to palbociclib but their growth was significantly inhibited by CDK4/6-D **(Fig. 2G)**. Similarly, although xenografts derived from T-47D cells overexpressing both CDK6-WT and CDK6-KD exhibited resistance to palbociclib, treatment with CDK4/6-D (5 mg/kg) significantly inhibited the growth of these xenografts. To assess target engagement of CDK4/6-D *in vivo*, we performed Western blot analysis for CDK2, 4, 6 and 9 in tumors harvested from CDK4/6-D-treated mice. We observed significant depletion in the levels of CDK4 and 6 in these tumors while there was no significant change in the protein levels of CDK2 and 9 relative to tumors from mice treated with vehicle. Our findings provide strong and direct evidence that kinase-independent function of CDK6 is sufficient to drive palbociclib resistance *in vivo* and that CDK4/6-D, by eliminating both the kinase and non-kinase functions of the protein, inhibits the growth of these tumors.

In summary, primary ER+ breast tumors show very low to undetectable expression of CDK6. Such CDK6 expression pattern, in combination with very low (<1%) frequency of *RB1* loss (*10*), makes them uniquely susceptible to CDK4/6 inhibitors, which are highly effective in blocking the kinase function of CDK4 (and 6). In response to treatment with CDK4/6 inhibitors, some ER+ BC tumors exhibit compensatory overexpression of CDK6, which enables them to exploit the non-kinase function of CDK6 to proliferate in the face of inhibition of kinase functions of CDK4 and 6. Our findings are consistent with a previous study which showed that a kinase-independent transcriptional function of CDK6 promotes tumor progression in acute lymphoid leukemia by promoting angiogenesis (*8*). In conclusion, we identify a novel mechanism of resistance to CDK4/6 inhibitors that relies on the non-kinase function of CDK6, thereby by enabling BC cells to achieve independence from the kinase functions of CDK4 and 6 to progress from G1 to S phase of cell cycle. Our findings not only challenge the current dogma that CDK4/6-mediated phosphorylation of pRB protein is necessary for cancer cells to progress from G1 to S phase of cell cycle but also highlight the limitations of the current therapeutic approaches that merely target the kinase function of CDK6 (and CDK4). Our study also provides a conceptual framework for understanding why ER+ breast tumors, which express very low to undetectable levels of CDK6, show remarkable clinical response to CDK4/6 inhibitors while most other cancers (which typically express high levels of CDK6) fail to derive clinical benefit from CDK4/6 inhibitors due to intrinsic resistance.

## Data Availability

All relevant data has been included in the manuscript. The reagents described in this manuscript may be requested through a Material Transfer Agreement.

## Acknowledgements

We acknowledge the assistance of the University of Texas Southwestern Tissue Management Shared Resource, a shared resource at the Simmons Comprehensive Cancer Center supported in part by the National Cancer Institute under award number P30 CA142543. We acknowledge the resources and services of the institutionally supported UT Southwestern Preclinical Pharmacology Core for conduct of pharmacokinetic studies. We acknowledge access to previously published breast cance patient-derived xenograft transcriptomic data provided by Dr. Alana L. Welm at Huntsman Cancer Institute, University of Utah, Salt Lake City, UT, USA. We acknowledge receipt of BSJ-1-184 (referred to as CDK4/6-D in this manuscript) from Dr. Nathanael Gray, Stanford University, CA, USA. This study was funded by grants to PGA from US Department of Defense (Breast Cancer Research Program Breakthrough Award W81XWH-21-1-0112), Charles Y. C. Pak Foundation and METAvivor.

## Author contributions

S.M.N.U., B.J., C.M.L., N.S.W and P.G.A. designed experiments. S.M.N.U., B.J., K.P., and N.S.W. performed the studies. S.M.N.U., B.J., C.M.L., S.S., N.S.W., P.G.A analyzed the data. S.M.N.U., B.J., C.M.L., N.S.W., and P.G.A. prepared figures and wrote the manuscript.

## Ethics Declarations

**Figure S1.**
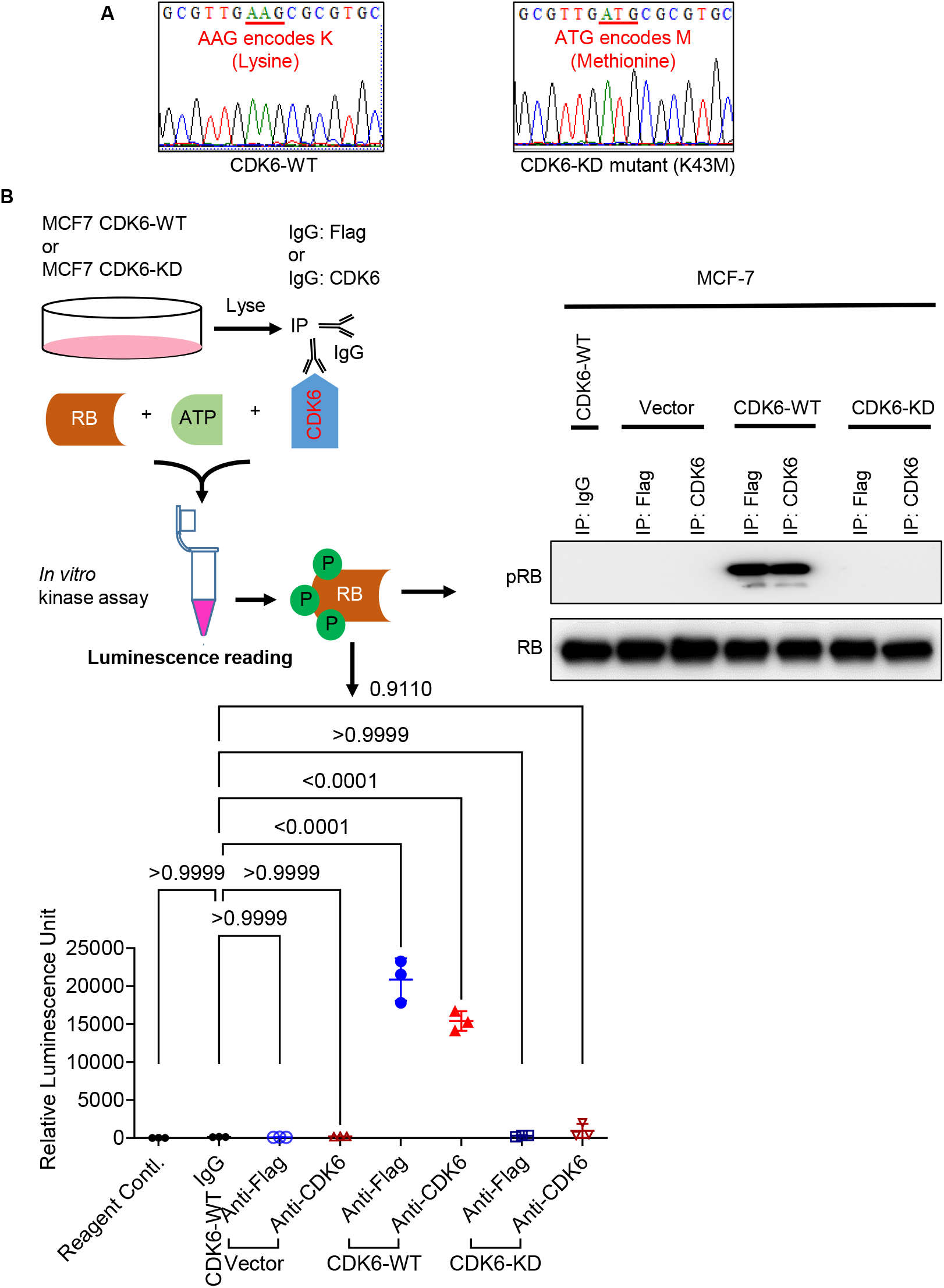
Functional characterization of a kinase-dead mutant (K43M) of CDK6. A: DNA sequencing confirms the presence of K43M mutation in CDK6-KD gene sequence. B: Schematic depiction of IP-*in vitro* kinase assay followed by ADP-Glo kinase assays and western blotting to evaluate the kinase function of CDK6-WT and CDK6-KD proteins. IP = immunoprecipitation; ADP = Adenosine diphosphate; WT= wild type; KD= kinase dead. Statistical significance was evaluated using ANOVA with Dunnett’s test.

**Figure S2.**
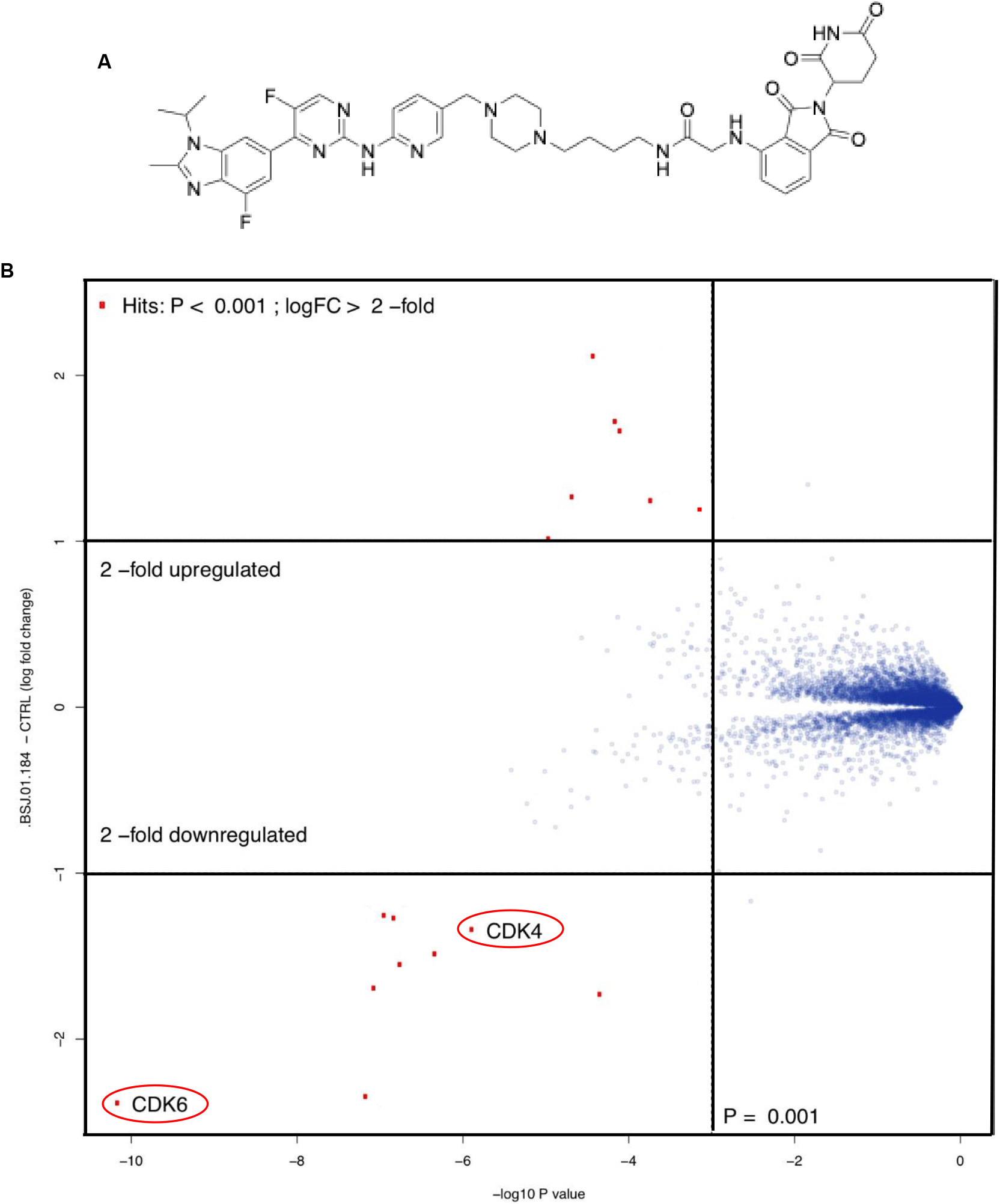
Proteome-wide selectivity of CDK4/6-D. A: Chemical structure of CDK4/6-D. B: Quantitative proteomics following treatment of Jurkat cells with 250 nM CDK4/6-D for 6 hours shows high selectivity for depletion of CDK6 (and CDK4) across the whole proteome.

**Figure S3.**
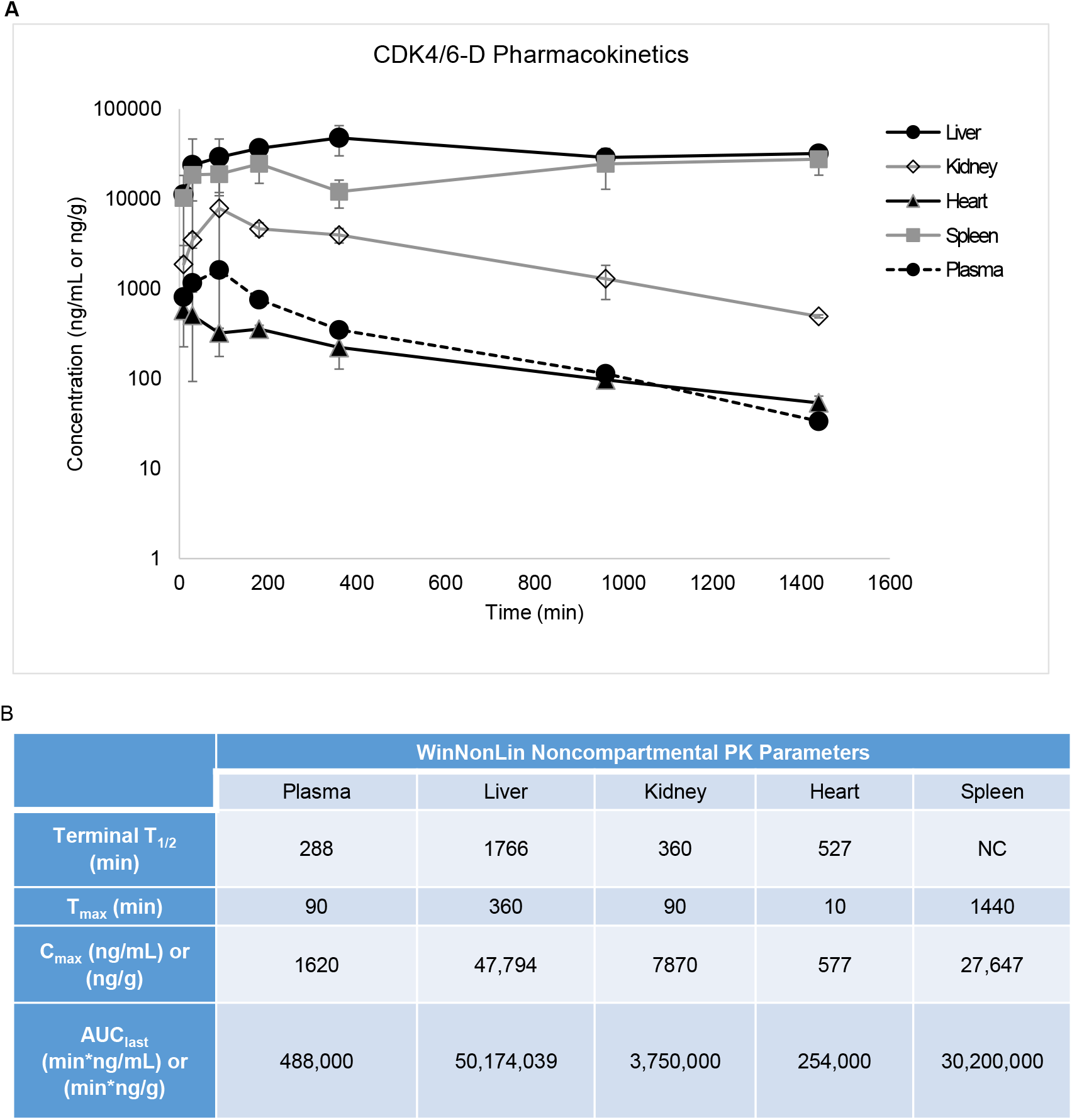
Plasma and tissue pharmacokinetics for CDK4/6-D. A-B: 6-week-old female CD1 mice were dosed by intraperitoneal injection in groups of three with 50 mg/kg CDK4/6-D, and blood and tissues were collected at various time points as indicated. After extraction of plasma and tissue homogenates with organic solvent, the supernatant was analyzed by liquid chromatography-tandem mass spectrometry. T_1/2_= Half life; T_max_= Time to maximum concentration; C_max_= Maximum drug concentration; AUC = Area Under the Curve

**Figure S4.**
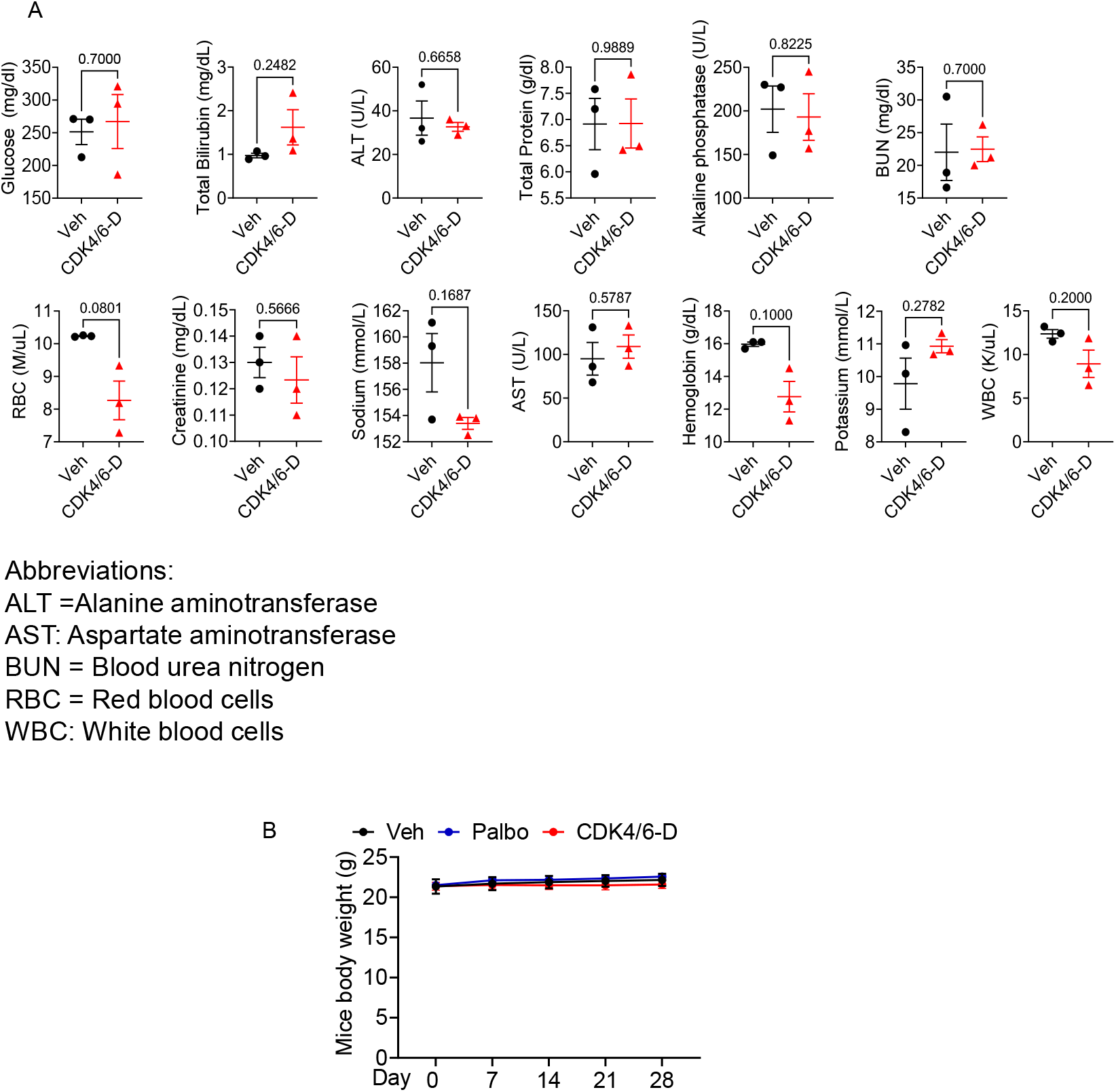
Toxicity studies in mice treated with CDK4/6-D. A: Blood collected from 8-week-old female C57BL/6 mice treated with CDK4/6-D (5 mg/kg by i.p. injection daily) for 1 week was subjected to biochemical analysis as indicated to assess the function of organs such as kidney, liver and bone marrow. Statistical significance was determined using unpaired, two-tailed t-test. ns = p > 0.05. B: Body weights of 8-week-old female C57BL/6 mice (n=5) treated with vehicle, palbociclcib (100 mg/kg by oral gavage daily) or 5 mg/kg CDK4/6-D (5 mg/kg by i.p. injection daily) for 4 weeks.

**Table ST1.**
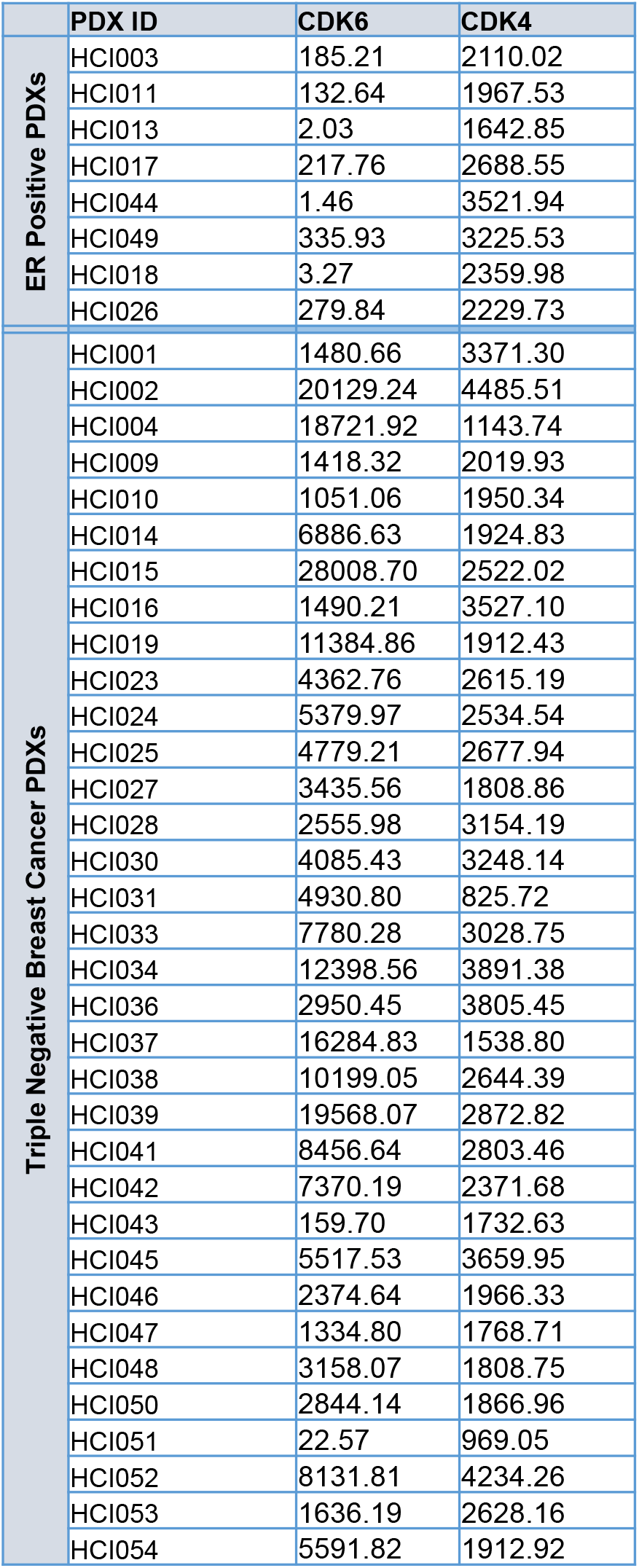
Raw data used for generation of Figure 1C showing normalized counts for CDK4 and CDK6 in the indicated PDXs. PDX= patient-derived xenograft.

