## Supplementary material for "The non-kinase function of CDK6 is a key driver of acquired resistance to CDK4/6 inhibitors in Estrogen Receptor-positive breast cancer": Methods

### Cell culture

CAMA1, MCF-7, MDA-MB-157, MDA-MB-231, MDA-MB-436 and MDA-MB-468 cell lines were obtained from the American Type Culture Collection (ATCC). HCC1428 was obtained from Hamon Cancer Center. T-47D and HCC1428 cells were maintained in Roswell Park Memorial Institute (RPMI) Medium (Gibco) with 10% Fetal Bovine Serum (FBS) from Hyclone, while CAMA1, MCF-7, MDA-MB-157, MDA-MB-231, MDA-MB-436, and MDA-MB-468 cells were cultured in Dulbecco's Modified Eagle Medium (DMEM) (Gibco) supplemented with 10% FBS. All cells were incubated in a humidified environment containing 5% $CO_2$. Cell viability was assessed and cell counts were performed using a Countess II FL Automated Cell Counter (Life Technologies, Gaithersburg, MD). Additionally, weekly screenings for mycoplasma contamination were conducted using MycoAlert Mycoplasma Detection Kit (Lonza Bioscience).

### Chemicals/Drugs

β-estradiol pellets (0.17 mg, 2-week release) were purchased from Innovative Research of America. Palbociclib was purchased from MedChemExpress. The synthesis and characterization of BSJ-1-184 (referred to as CDK4/6-D in this manuscript) has been previously described (*1*).

### Plasmids and Viral Transduction

The Lentiviral expression constructs encoding wild-type human CDK6 (CDK6 WT; pLV[Exp]-Puro-CMV>hCDK6[NM_001145306.2]/FLAG, Vector ID: VB231011-1521pau) and the kinase-dead (K43M) mutant of CDK6 (pLV[Exp]-Puro-CMV>hCDK6[NM_001145306.2]*K43M/FLAG, Vector ID: VB231011-1515jmm) were obtained from VectorBuilder.

**Generation of stable cell lines overexpressing CDK6**

Lentiviral particles were generated by co-transfecting HEK293T cells with the respective CDK6 constructs and standard packaging (psPAX2) and envelope (pMD2.G) plasmids using Lipofectamine 3000 according to the manufacturer's protocol (ThermoFisher Scientific). Viral supernatants were harvested 12-48 hours post-transfection, filtered, and used to transduce MCF-7 and T-47D cells in the presence of polybrene (8 µg/mL) (Millipore). Puromycin (InvivoGen, San Diego, USA) selection was used to generate stable cell lines expressing the appropriate expression vector.

**Cell growth assays**

Cells were seeded in a 6-well tissue culture plate in DMEM or RPMI medium supplemented with 10% FBS, as indicated, and treated with indicated drugs 24 hours later. The cells were incubated for 10 days, with media and drug changes every three days. On day 10, the cells were washed, fixed with 4% paraformaldehyde, and stained with 0.5% crystal violet solution for 20 minutes. After washing with water, the plates were air-dried, and photographs were taken. The crystal violet was then solubilized using 20% acetic acid, and absorbance was measured at 490 nm using a Spark 10M multimode microplate reader (Tecan).

**Cell cycle analysis**

Cells seeded in 35 mm dish were treated with the indicated drug and incubated for 24 hours. Cells were then trypsinized, pelleted at 1,000 rpm for 5 minutes, resuspended in 500 µL phosphate-buffered saline (PBS), and fixed by dropwise addition of pre-chilled 70% ethanol with gentle vortexing. Fixation was carried out on ice for 1 hour. Fixed cells were centrifuged at 2,000 rpm for 5 min, the supernatant was removed leaving ~100 µL residual ethanol, and cells were resuspended by vortexing. For propidium iodide (PI) staining, cells were incubated in 500 µL of staining buffer (PBS containing 0.1% Triton X-100, 40 µg/mL RNase A, and 2 µg/mL PI) at 37°C for 15–20 minutes in the dark. Samples were filtered through a nylon mesh to remove clumps. The proportion of cells in each phase of the cell cycle (G0/G1, S and G2/M) was determined by flow cytometry based on DNA content.

**Immunoblotting**

Tumor tissues were homogenized using the FastPrep-24™ 5G Homogenizer (MP Biomedicals) with Lysing Matrix S steel beads (Cat. #6925000, MP Biomedicals) in a modified Radioimmunoprecipitation Assay (RIPA) lysis buffer containing 50 mM Tris-HCl (pH 7.5), 1% NP-40, 0.25% sodium deoxycholate, 150 mM sodium chloride, 1 mM ethylenediaminetetraacetic acid (EDTA), and complete protease and phosphatase inhibitor cocktails (Roche). Tumor cells were lysed using the same lysis buffer in the culture plate and protein lysates were collected following centrifugation. Equal amounts of protein were resolved by Sodium Dodecyl Sulfate (SDS)-Polyacrylamide Gel Electrophoresis (PAGE) and transferred onto polyvinylidene fluoride (PVDF) membranes. The membranes were immunoblotted with the indicated antibodies.

CDK2 (Cat. # 18048), CDK4 (Cat. # 12790), CDK6 (Cat. # 3136), CDK9 (Cat. # 2316), Rb (Cat. #9309), Phospho-Rb Ser807/811 (Cat. #8516), ERα (Cat. # 8644), and β-actin (Cat. #3700), all from Cell Signaling Technology, and anti-FLAG (Cat. # F3165) from Sigma-Aldrich. Immunoreactive proteins were detected using ECL Super Signal West Femto substrate reagent (ThermoFisher Scientific).

**Immunohistochemistry (IHC)**

Formalin-Fixed Paraffin-Embedded slides were baked for 20 minutes at 60$^{o}$C, then deparaffinized and hydrated before the antigen retrieval step. Heat-induced antigen retrieval was performed at pH 9.0 for 20 minutes. The tissue was incubated with a peroxidase block followed by CDK4 antibody, 1:50 dilution (mouse anti- CDK4; cat# sc-23896; Santa Cruz Biotechnology) or CDK6 antibody, 1:250 dilution (mouse anti-CDK6 cat# sc-7961; Santa Cruz Biotechnology). A boardcertified breast pathologist (SS) reviewed the slides and categorized them as detectable or undetectable for expression of CDK4 and CDK6.

**Immunoprecipitation (IP) - *in vitro* kinase Assay**

The IP-*in vitro* kinase assay to quantify kinase activity of CDK6-WT and CDK6-KD proteins was carried out as previously described (*2*).

**Proteome-wide selectivity of CDK4/6-D**

The proteome-wide selectivity of CDK4/6-D was assessed in Jurkat cells, which endogenously express high levels of CDK4 and CDK6. Following treatment with 250 nM CDK4/6-D or vehicle for 6 hours, we performed multiplexed mass spectrometry (MS)-based proteomic analysis as previously described (*3*).

**Analytical Liquid Chromatography-Tandem Mass Spectrometry (LC-MS/MS)**

An LC-MS/MS assay using an AB Sciex (Framingham, MA) 4000 QTRAP® mass spectrometer coupled to a Shimadzu (Columbia, MD) Nexera LC for quantifying the plasma and tissue levels of CDK4/6-D. The compound was detected with the mass spectrometer in positive Electrospray Ionization - Multiple Reaction Monitoring (ESI-MRM) mode by following the precursor to fragment ion transition 863.380 to 393.100. An Agilent C18 XDB column (5 micron, 50 x 4.6 mm) was used for chromatography of the compound with the following conditions: Flow rate 1.5 ml/min; Buffer A: dH20 + 0.1% formic acid, Buffer B: Methanol + 0.1% formic acid; 0 – 1.5 min 5% B, 1.5-2.0 min gradient to 100% B, 2.0 – 3.5 min 100% B, 3.5 – 3.6 gradient to 5% B, 3.6 - 4.6 5%B. Palbociclib (transition 448.097 to 380.036) from Medkoo Biosciences (Durham, NC) was used as an internal standard (IS).

**Pharmacokinetic (PK) studies**

PK studies were performed by dosing 6-week old CD1 female mice (Charles River, Willmington, MA) with CDK4/6-D by an intraperitoneal route at 50 mg/kg, 0.2 mL/mouse, formulated in 5% DMSO/95% of a 10% solution of captisol in water. Animals were sacrificed in groups of three, blood was obtained by cardiac puncture at each time point (0, 10, 30, 90, 180, 360, 960 and 1440 minutes post dose) using the anticoagulant, acidified citrate dextrose (ACD) and plasma isolated by centrifugation for 10 minutes at 9600 x g. Tissues were harvested, rinsed with PBS and blotted dry, weighed and snap frozen. Tissues were subsequently homogenized in a 3X volume of PBS by weight using a polytron homogenizer. 100 µl of plasma or tissue homogenate was mixed with a 2X volume of methanol containing formic acid (0.15%) and 37.5 ng/mL palbociclib as an internal standard.

The samples were vortexed for 15 seconds, incubated at room temp for 10 minutes and spun twice at 16,100 x g in a refrigerated (4ºC) microcentrifuge. The resulting supernatants were evaluated by LC-MS/MS as described above. Standard curves were generated using blank plasma (Bioreclamation, Westbury, NY) or blank tissue homogenate spiked with known concentrations of compound and processed as described above. The concentrations of drug in each time-point sample were quantified using Analyst software (Sciex). Compound in tissue vasculature was subtracted prior to plotting and the concentration of each compound measured in plasma. Compounds were assumed to partition equally between red blood cells and plasma. A value of 3-fold above the signal obtained from blank plasma was designated the limit of detection (LOD). The limit of quantitation (LOQ) was defined as the lowest concentration at which back calculation yielded a concentration within 20% of theoretical. Pharmacokinetic parameters were calculated using the noncompartmental analysis tool of Phoenix WinNonLin (Certara/Pharsight, Sunnyvale, CA).

***In vivo* efficacy studies:**

10 million MCF-7 or T-47D cells were suspended in a 1:1 mixture of Matrigel and DMEM and injected subcutaneously into the flanks of 6 to 8-week-old ovariectomized female nude mice (Charles River). A β-estradiol pellet (0.17 mg, 2-week release) was subcutaneously implanted in the neck area to support tumor growth. Once tumors reached 100-150 $mm^3$ in size, mice were randomized to the following treatments groups: Vehicle by oral gavage daily, 6 days per week; Palbociclib (100 mg/kg) by oral gavage daily, 6 days per week; and CDK4/6-D (5 mg/kg) by intraperitoneal injection daily, 6 days per week. Mice were sacrificed 4 weeks later and tumors were harvested for pharmacodynamic studies. All animal experiments were conducted in accordance with the UT Southwestern Institutional Animal Care and Use Committee (IACUC) approved protocol.

**Statistics**

*In vitro* studies were performed in biological triplicates as indicated. Statistical analyses were conducted using GraphPad Prism version 10 (GraphPad Software). Pairwise comparisons between groups were made using a two-tailed Student's t-test. For analyses involving more than two groups, one-way ANOVA followed by Dunnett's test was used to compare each experimental group to the control. A p value of less than 0.05 was considered statistically significant.
